# Major effect mutations drive DNA methylation variation after colonization of a novel habitat

**DOI:** 10.1101/2025.06.14.659694

**Authors:** Johan Zicola, Emmanuel Tergemina, Mehmet Göktay, Célia Neto, Robert J. Schmitz, Angela M. Hancock

## Abstract

DNA methylation is important to maintain genome stability, but alterations in genome-wide methylation patterns can produce widespread genomic effects, which have the potential to facilitate rapid adaptation. We investigate DNA methylation evolution in *Arabidopsis thaliana* during its colonization of the drought-prone Cape Verde Islands (CVI). We identified three high impact changes in genes linking histone modification to DNA methylation that underlie variation in DNA methylation within CVI. Gene body methylation is reduced in CVI relative to the Moroccan outgroup due to a 2.7-kb deletion between two *VARIANT IN METHYLATION genes* (*VIM2* and *VIM4*) that causes aberrant expression of the *VIM2/4* homologs. Disruptions of *CHROMOMETHYLASE 2 (CMT2)* and a newly identified DNA methylation modulator, *F-BOX PROTEIN 5 (FBX5),* which we validated using CRISPR mutant analysis, contribute to DNA methylation of transposable elements (TEs) within CVI. Overall, our results reveal rapid methylome evolution driven largely by high impact variants in three genes.

## Main

Epigenetic modifications of DNA, primarily 5-methylcytosine, are crucial for maintaining genome stability^1–4^. While methylation in mammals occurs primarily at CG dinucleotides, plants exhibit methylation in all cytosine contexts: symmetric CG and CHG, and asymmetric CHH (where H is C, A, or T). DNA methylation serves multiple functions, including silencing transposable elements (TEs), regulating gene expression, processing RNA, and mediating chromosome interactions^4,5^.

In plants, distinct methyltransferases govern different methylation contexts: METHYLTRANSFERASE 1 (MET1) maintains CG methylation, while CHROMOMETHYLASE 3 (CMT3) maintains CHG methylation^6–12^. CHH methylation is established by DOMAIN REARRANGED METHYLTRANSFERASE 1/2 (DRM1/2) through the RNA-directed DNA methylation (RdDM) pathway^13,14^, whereas CHROMOMETHYLASE 2 (CMT2) acts in heterochromatic regions independently of small RNA activity^12,15^. A feedback loop exists between histone modifications and DNA methylation, exemplified by CMT3’s dependence histone 3 lysine 9 dimethylation (H3K9me2) deposited by SU(VAR)3-9 HOMOLOG 4 (SUVH4) (also called KRYPTONITE)^8,9,9,11,16–19^. Additionally, small euchromatic TEs are targeted by RdDM through siRNAs and maintained by DRM2, while methylation of long heterochromatic TEs requires DECREASED DNA METHYLATION 1 (DDM1) in conjunction with CMT2^15^.

In *Arabidopsis thaliana* (Arabidopsis), genome-wide loss of DNA methylation affects gene expression and TEs repression, but it is not lethal^10,15,20,21^. While CG-dinucleotide methylation of promoters typically suppresses transcription^22^, it is relatively rare in plants^23^. Gene body methylation (gbM), predominately in the CG context, occurs in approximately 20% of Arabidopsis genes and correlates with moderate and constitutive expression^24–34^. GbM is maintained by MET1 and likely also requires CMT3 for its initiation and maintenance since plant species without *CMT3* lack gbM^35,36^. Despite ongoing debate about whether gbM is merely a byproduct of TE silencing^35,37,38^, or serves functions such as gene expression stabilization, cryptic transcription inhibition, and protection against TE insertion^39,40^, conservation across diverse plant species suggests functional importance^30,39,41,42^.

Studying natural variation in DNA methylation can provide insights into how epigenetic changes evolve over time in natural populations and connect to other physiological and molecular changes^31,43–45^. Previous studies of natural variation in gbM among Arabidopsis accessions identified Cvi-0, from the Cape Verde archipelago, as having the lowest gbM levels^43,46,45,47^. Historical reconstructions indicate that Arabidopsis colonized this drought-prone region approximately 5,000 years ago from Morocco through a strong bottleneck, that eliminated most standing genetic variation, necessitating rapid adaptation through new variants^48^.

Here, we investigate DNA methylation patterns in the CVI population, hypothesizing that mutations affecting genome-wide epigenetic profiles could influence many genes simultaneously, facilitating rapid adaptation. We demonstrate reduced mean gbM in this population compared to both the closest Moroccan outgroup and well-studied Eurasian populations. Through genome-wide association studies of methylation signatures across different cytosine contexts, we identify novel alleles in known methylation modulators (VIM2/4 and CMT2) and discover a previously uncharacterized methylation regulator (FBX5), with effects confirmed through CRISPR mutant analysis.

## Results

### Gene body methylation is reduced in Arabidopsis from Cape Verde

We assessed gbM variation in the Santo Antão population of 83 accessions from CVI using whole-genome bisulfite sequencing. For comparison, we also conducted whole-genome bisulfite sequencing on 22 representative accessions from Morocco, the closest outgroup population to CVI^48^ (**Fig. 1a)** and contrasted these to published whole-genome bisulfite sequencing data from Eurasian samples^45^ (**Fig. 1a**). Average gbM levels of CVI plants were significantly lower than in any other population (Wilcoxon-test, p-value < 2.2 x 10^-16^), including the Moroccan outgroup (Wilcoxon-test, p-value = 2.6 x 10^-11^), which exhibits gbM levels similar to other worldwide populations (**Fig. 1b**). We estimated SNP-based heritability^49^ of gbM levels in the CVI population to be 97% based on the correlation between phenotypic and genetic covariance in a Bayesian sparse linear mixed model that accounts for small and large effect variants^50^.

**Fig. 1.**
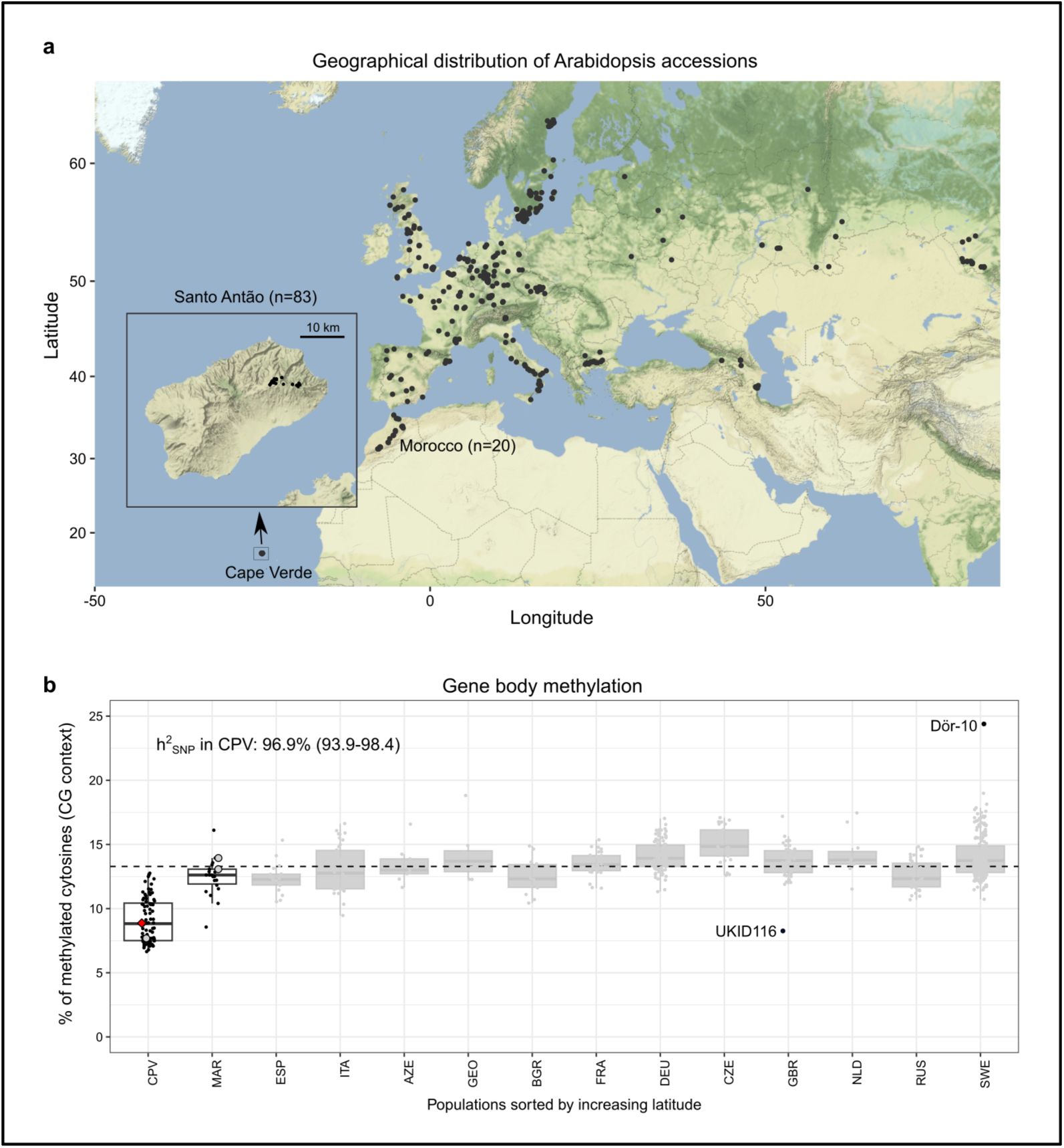
Arabidopsis populations from CVI diverge epigenetically from all other known populations. **a**, Geographical distribution of the bisulfite-sequenced accessions from the 1001GP and this study. Only countries with at least eight bisulfite-sequenced accessions are shown. New data were collected in this study for CVI and Morocco, indicated on the map and the inset. **b**, Gene body methylation for the CG context for the accessions shown in **a**, grouped by country (3-letters ISO names, CPV=CVI, MAR=Morocco), and ordered by increasing latitude. Center lines of boxplots show the medians, boxes denote the 25th and 75th percentiles, whiskers extend 1.5 times the interquartile range from the 25th and 75th percentiles, and dots represent individual data points. Dashed lines indicate the median value across all countries. Grey boxplots represent methylation levels from published data (representative subset of 523 accessions of the 1001GP). The red diamond shape indicates the methylation levels of Cvi-0 from our data. Grey circles indicate methylation levels for published data for Cvi-0 and MAR (two accessions). The highlighted accessions UKID116 (GBR) and Dör-10 (SWE) were previously identified as having the lowest and highest gbM levels worldwide, respectively^45^. SNP heritability (h^2^_snp_) for gbM in CPV and its 95% confidence interval are indicated on the plot.

### A 2.7 kb deletion of the *VIM2* and *VIM4* promoters explains the reduced gene body methylation in Cape Verde

To identify genetic modifiers of gbM in the CVI population, we conducted a genome wide association study (GWAS) of DNA methylome data using percent DNA methylation over gene bodies as a phenotype. We identified a major peak on chromosome 1 (**Fig. 2a**), although no SNP in that genomic region tagged a gene known to be involved in DNA methylation. This could occur if the peak was caused by an untyped genetic variant, such as a structural variant. To investigate the possibility that structural variation was responsible for the peak, we examined associations with k-mers^51^. To localize k-mers to the genome, we aligned significantly associated k-mers to the Col-0 reference assembly as well as to four newly created long read-based genome assemblies from CVI accessions (S1-1, S15-3, S5-10 and Cvi-0). A group of 1,657 of the most significant k-mers flanked the promoter regions of *VARIANT IN METHYLATION 2* (*VIM2*, AT1G66050) and *VIM4* (AT1G66040), which are positioned head-to-head approximately 2.7 kb apart in Col-0 (**Fig. 2b**). This variant is an excellent candidate because VIM2 and VIM4 are involved in CG methylation maintenance^52,53^. These k-mers were mappable in the S1-1 and S15-3 accessions, but unmappable in the Cvi-0 and S5-10 assemblies, consistent with a segregating deletion in Cape Verde. Based on Col-0 annotations, the full deletion contains three RC/helitron fragments and a reverse transcriptase-like gene (AT1G66045), suggesting that the deletion may have resulted from a transposition event.

**Fig. 2.**
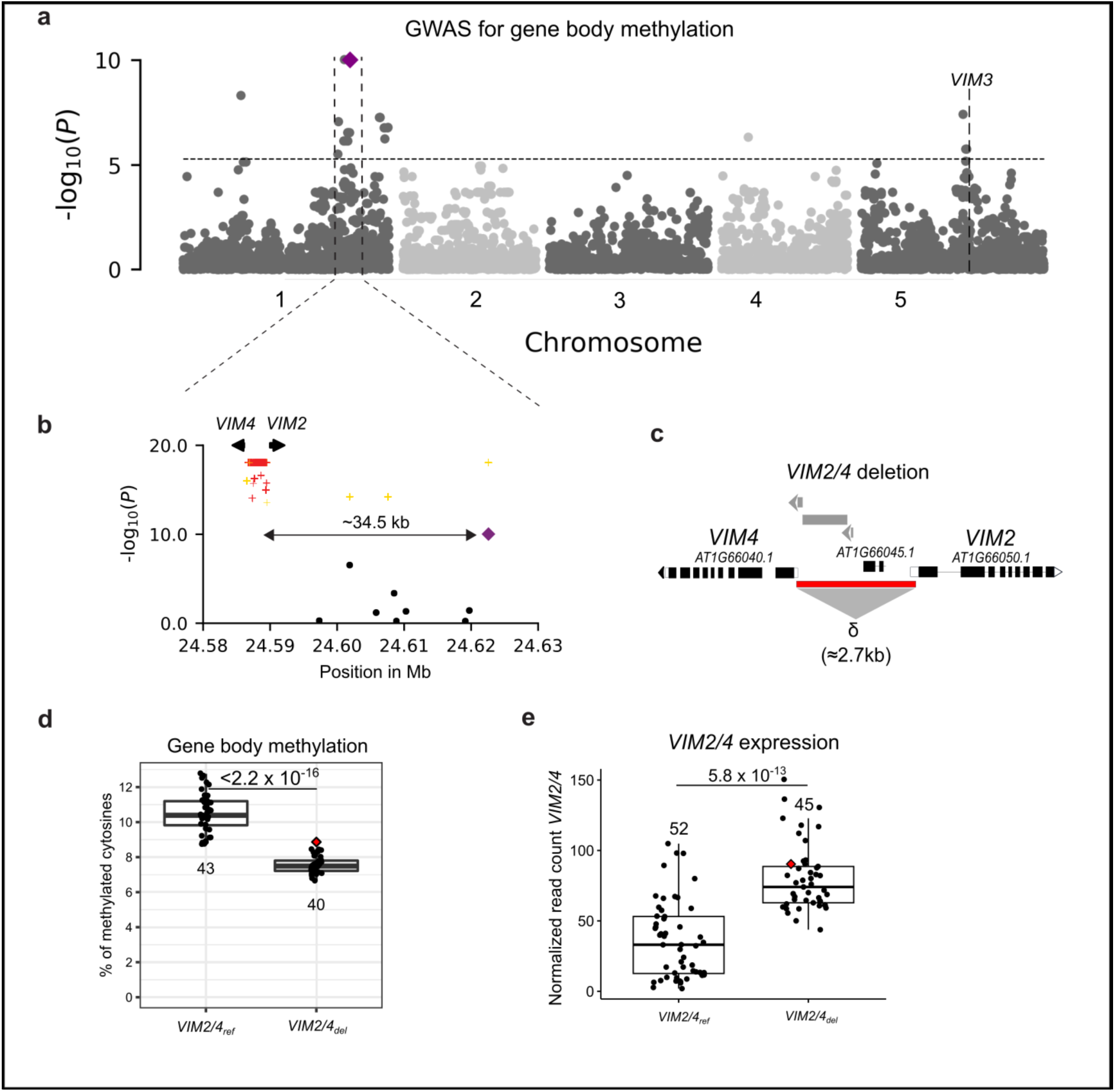
Loss of *VIM2* and *VIM4* promoter is associated with low gbM. **a**, Manhattan plot for the gbM GWAS in CVI. 10,321 SNPs were analyzed with a minor allele frequency (MAF) threshold of 5%. Bonferroni threshold is indicated by the dashed line. **b**, *k*-mer GWAS at the chromosome 1 peak. *k*-mers are highlighted in yellow if they mapped to the Cvi-0 assembly and in red if they did not map to the Cvi-0 assembly. SNPs are highlighted in black. The most significant SNP is annotated with a purple diamond. **c**, The deletion at *VIM2* and *VIM4* (yellow box) overlaps the 5’-UTRs (white boxes) without affecting the start codons. Grey boxes indicate the locations of TEs. **d**, Difference in gbM between the accession without (0) and with the *VIM2/4* deletion (1). A two-tailed Welch’s t-test was performed to compare the means of the groups. **e**, Normalized read count for *VIM2/4* expression levels in 97 accessions from CVI. A two-tailed t-test was performed between the two groups. In **d** and **e**, center lines of boxplots show the medians, box limits the 25th and 75th percentiles, whiskers extend 1.5 times the interquartile range from the 25th and 75th percentiles, and dots represent individual data points. Red diamond represents gbM for Cvi-0. Number of individuals for each group is indicated below each box. P-values are indicated on the plots.

By comparing the *VIM2/4* genomic region of the two CVI accessions carrying the deletion to the Col-0 reference assembly, we found that the start codons of the two *VIM* genes are preserved, but a part of the 5’UTR is removed for each gene, which could impact the regulation of *VIM2*/*VIM4* (**Fig. 2c)**. We also found that the CVI population carries a premature stop codon at amino acid position 563 of VIM4, indicating that VIM4 is probably nonfunctional in CVI. In addition to the peak at *VIM2/4*, we found a peak on chromosome 5 with the most significant SNP (chr5:15,047,549) within a TE (AT5TE54335) in the 4^th^ exon of a NOD26-like intrinsic protein (AT5G37810). This SNP is 790 kb upstream of the candidate gene *VIM3*, but we could not identify any potential SNP or SV that could affect *VIM3* function.

We genotyped the *VIM2/4* deletion in the CVI population and found that mean gbM levels were reduced by 27.9% for accessions carrying the *VIM2/4* deletion (*VIM2/4*_del_) relative to the reference allele (*VIM2/4*_ref_), corresponding to an absolute gbM reduction of 2.94% methylation in the linear mixed model (**Fig. 2d**). *VIM2/4*_del_ is found in 20 of the 24 sites where Arabidopsis was collected across Santo Antão, with an overall frequency of 47% (89/189) on the island. Using *VIM2/4*_del_ as a covariate, we did not identify additional regions significantly associated with variation in gbM, indicating that *VIM2/4*_del_ is sufficient to explain the variation in the region.

To assess the potential impacts of *VIM2/4*_del_ more broadly, we examined which regions were differentially methylated between accessions with the *VIM2/4*_ref_ and the *VIM2/4*_del_ allele. Using a threshold of 25% difference in mCG, we identified 12,996 differentially methylated regions (DMRs) with 94% showing lower mCG levels in the accessions with *VIM2/4*_del_. Most of the DMRs (92%, n=11,948) overlap genic regions (n=7,541), while 8% (n=978) overlap TEs. Out of the 27,655 protein-coding genes from the Araport11 annotation, 26% (n=7,200) co-localize with at least one DMR.

GO enrichment analysis on protein-coding genes overlapping DMRs revealed a strong enrichment for the cellular components ontology GO term Cul4-RING E3 ubiquitin ligase complex (GO:0080008; 2.4 fold, Benjamini-corrected p-value = 2x10^-15^), which was driven by 79 genes, many of which are known to be involved in environmental response traits, including light signaling, flowering, and DNA repair. Biological processes enriched for DMRs included those directly related to expected large-scale effects of DNA methylation, such as regulation of gene expression (1.8-fold, Benjamini-corrected p-value = 8x10^-57^) and chromatin remodeling (2.6-fold, Benjamini-corrected p-value = 5.9x10^-10^), as well as biological processes related to DNA repair (2.3-fold, Benjamini-corrected p-value = 2.4x10^-22^) and embryo development ending in seed dormancy (1.8-fold, Benjamini-corrected p-value = 2.4x10^-30^).

Leaf transcriptomes clustered mainly by population, which is expected if transcriptome variation is due to genetic factors. Ninety-four genes were significantly differentially expressed between the *VIM2/4*_del_ and *VIM2/4*_ref_ genotypes. Converse to our expectations, expression of *VIM2* and *VIM4* was increased in *VIM2/4*_del_ accessions. While we expected the loss of the promoter regions in *VIM2/4*_del_ accessions would abolish their expression, we actually found that the genes were upregulated when the UTRs were deleted, suggesting that they still have functional minimal promoters and the UTR contains repressors (**Fig. 2e)**.

Although it has been shown that the triple *vim1/2/3* triple mutant leads to a strong reduction in mCG^53–55^, the mCG levels are higher in the *vim2* single mutant relative to wild type, while mCG levels are reduced in *vim1*, *vim3*, and *vim1/2/3*. This observation suggests that the function of VIM2 is more complex and that it could actually be a negative regulator of mCG, which would explain why overexpression of *VIM2/4* in the accessions with *VIM2/4del* is associated with gbM reduction.

Gene ontology (GO) term enrichment analysis on the 94 DEGs showed significant enrichment only for molecular functions related to methylcytosine binding (GO:0010429, GO:0008327, GO:0010428), which was due to the three *VIM* homologs (*VIM2*, *VIM3* and *VIM4*) present in the DEG list (>100-fold enrichment, Benjamini-corrected p-value < 3.8x10^-2^). Interestingly, a DMR overlaps the second exon of *VIM3,* with a lower CG methylation in VIM2/4_del_ genotype, which could potentially be responsible for the covariance in *VIM3* expression with the *VIM2*/*VIM4* deletion. The fact that we do not see evidence of a change in gene expression in the categories for which we see a change in methylation is consistent with evidence that gene body methylation is not directly linked to gene expression^37^, but may instead be linked to moderate stochastic changes over longer time scales.

### Transposable element DNA methylation in Cape Verde Arabidopsis is highly heritable

Next, we examined DNA methylation over TEs in CVI. We first assessed heritability of DNA methylation variation at all annotated TEs (31,189) and found SNP heritability to be 70-80% (**Fig. 3a-c**). We then considered separately long TEs (>4kb), which are methylated specifically by the DDM1-CMT2 pathway^15^ so that methylation of these TEs may have a less complex genetic architecture. The 1,235 long TEs retained are mainly comprised of Class I long terminal repeat (LTR) Gypsy (42%) and Copia (17%) TEs, and of Class II DNA MuDR TEs (16%). Long TEs are about twice as highly methylated as TEs overall and their methylation is more highly heritable in the CG and CHH contexts. Phenotypic variance was highest for mCHG and mCHH-methylated long TEs (**Fig. 3d-f**). SNP heritability was highest for mCHH (94.3%), followed by mCG (81.2%) in long TEs (**Fig. 3c, e**). For mCHG, the SNP heritability was higher when considering all TEs (72.4%) than long TEs (59.5%). Overall, the high heritability of TE methylation indicates strong genetic effects and suitability for mapping using GWAS. In subsequent analyses we focus on long TEs because methylation in these contexts is highly heritable.

**Fig. 3.**
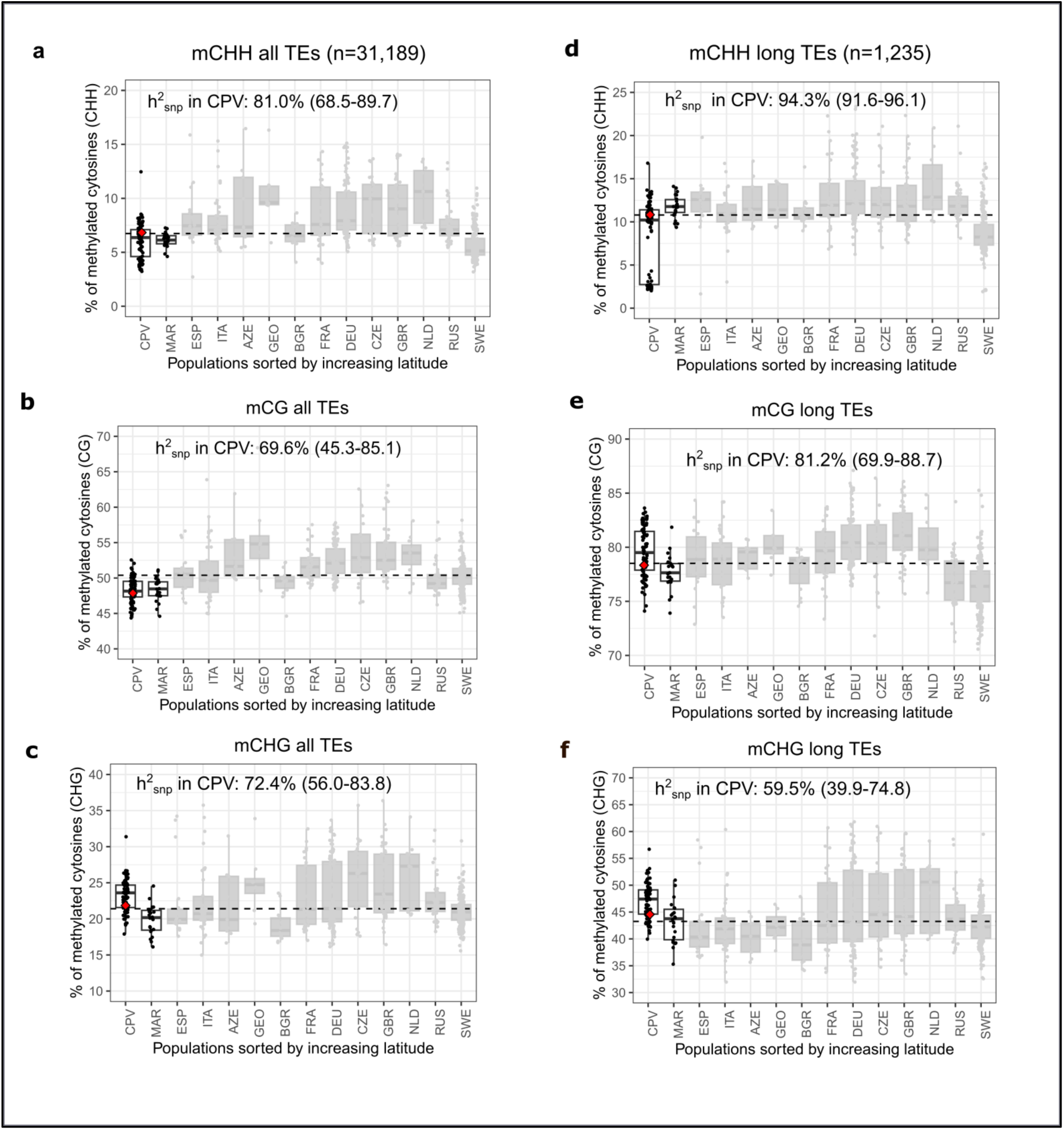
DNA methylation at TEs shows a large variance in CVI. DNA methylation in CHH, CG, and CHG contexts for all TEs (**a-c**) and long TEs (>4 kb) (**d-f**). Accessions grouped by country (3-letter ISO names, CPV=CVI, MAR=Morocco), and ordered by increasing latitude. Dashed lines indicate the median value across all countries. Grey boxplots represent the methylation levels from data (subset of 523 accessions of the 1001GP). The red diamond shape indicates the methylation levels of Cvi-0.

### *CMT2* loss of function reduces mCHH over long TEs

GWAS using mCHH over long TEs as a phenotype revealed a clear peak on chromosome 4 (**Fig. 4a**), where the most significant SNP is a nonsense mutation in *CMT2* that introduces a stop codon and truncates the protein before a C-terminal methyltransferase domain (Chr4:10,420,088, A>T, **Fig. 4b**). Note that Cvi-0 has the *CMT2_ref_* allele indicating that the gene is functional in this accession (**Fig. 4c**). *CMT2_stop_* is associated with a reduction of mCHH over long TEs of ≈82% (β coefficient = -4.18). *CMT2_stop_* is present at 14 geographic sites with an overall frequency of 31% in Santo Antão (60 out 189 accessions). Within those 14 sites, the allele segregates in 13 of the 14 sites, and the only exception is a site with a very small sample size (n=4). This pattern suggests the allele may be subject to balancing selection due to a tradeoff between its negative evolutionary effects through transposon mobilization and its potential positive effects due to increased heat tolerance of plants carrying the allele^44,56^.

**Fig. 4.**
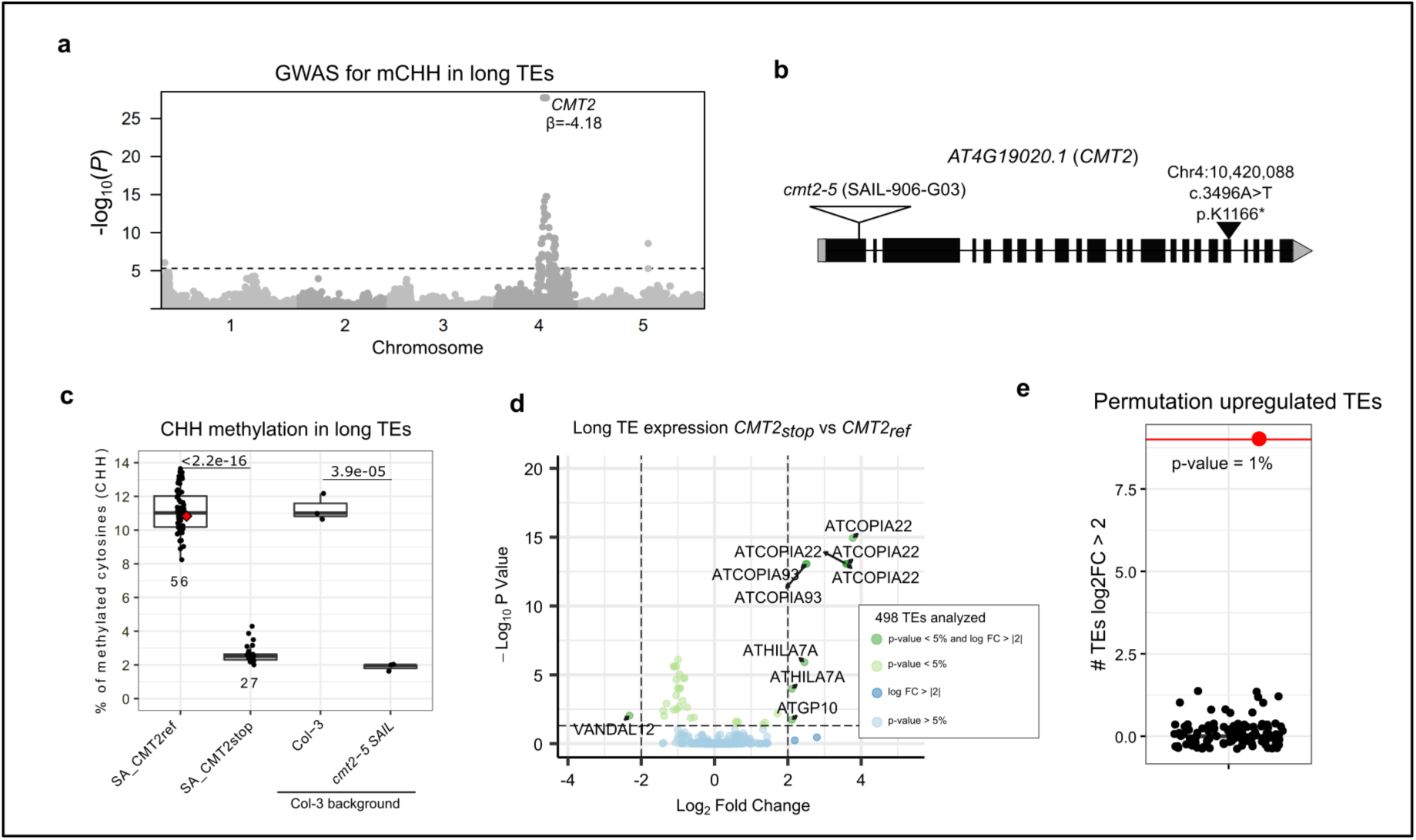
Nonsense variant in *CMT2* explains most of mCHH variation in long TEs. **a**, Manhattan plot of GWAS for mCHH in in long TEs (>4 kb). The nonsense SNP at CMT2 (Chr4:10,420,088) is highlighted in blue. Bonferroni threshold is indicated by a dashed line. **b,** Representation of the transcript model *AT4G19020.1* of *CMT2* includes exons (black boxes), UTRs (grey boxes), the location and HGVS nomenclature information of the nonsense variant located in the 19^th^ exon (black triangle), and the T-DNA insertion of *cmt2-5* SAIL mutant (white triangle). **c,** Levels of mCHH in long TEs for Santo Antão (SA) individuals with the reference allele (*CMT2_ref_*) and the nonsense allele (*CMT2_stop_*), and for the individuals with the T-DNA insertion line *cmt2-5* (SAIL_906_G03) and the Col-3 wild-type background accession. Red diamond represents mCHH level for Cvi-0. Number of individuals for each group is indicated below each box for Santo Antão accessions, three biological replicates shown for Col-3 and *cmt2-5*. Two-tailed t-tests were performed to compare the means of the two groups (p-value indicated on the plot). Center lines of boxplots show the medians, box limits the 25^th^ and 75^th^ percentiles, whiskers extend 1.5 times the interquartile range from the 25^th^ and 75^th^ percentiles, dots represent individual data points. **d,** Volcano plot of long TE expression in accessions with the *CMT2_stop_* vs *CMT2_ref_* alleles. TEs with significant differential expression (< 5% adjusted p-value) and a log2 fold change > |2| are displayed on the plot. Circles indicate TEs belonging to the same family. NS stands for non-significant. 524 TEs had at least 10 reads across all accessions and could be used for differential expression analysis. **e**, Number of significantly upregulated TEs (log2FC > 2) based on 100 permutations (black dots) with the observed values indicated in red (9 TEs). P-value is indicated on the plot and calculated by using the formula (B+1)/(M+1), where B is the number of permutations in which a value greater or equal than the observed value is obtained and M is the total number of permutations sampled.

An additional peak in chromosome 5 contains two SNPs located in a TE (AT5TE52400) (**Fig. 4a**) that maps 472-749 kb downstream of *CMT2* on chromosome 4 in the Cape Verde genome assemblies. Therefore, the peak at chromosome 5 is an artifact of a discrepancy between the reference genome assembly (Col-0 TAIR10) used for mapping during SNP calling and the CVI genome organization. We did not find other significant peaks when *CMT2_stop_* was used as a covariate. Comparison of CHH methylation in the *CMT2stop* and *CMT2_ref_* alleles in the natural population to a *cmt2-5* T-DNA insertion mutant, where mCHH was similar, provided further evidence that a truncation of CMT2 could explain the observed patterns. We also compared the difference mCHH in *CMT2* alleles in the natural population to a *cmt2-5* T-DNA insertion mutant and its WT background accession Col-0 and found the change in mCHH was similar, indicating that the natural *CMT2_stop_* allele in CVI is most likely a null mutant (**Fig. 4c**). Overall, our data indicate that most of the variation in mCHH in CVI is driven by the natural *CMT2_stop_* allele.

Next, we tested whether changes in mCHH due to the *CMT2stop* allele could affect long TE expression and found 63 significantly differentially expressed TEs, including 23 downregulated and 15 upregulated (**Fig. 4d**). The upregulated TEs are enriched in the LTR/Copia superfamily, with the families ATCOPIA22 and ATCOPIA93 showing the most significant upregulation (**Fig. 4d**). We tested whether this expression pattern was likely to be found by chance by conducting 100 permutations in which we attributed randomly chosen *CMT2* alleles within CVI subpopulation and found that the number of significantly differentially expressed TEs was always well above the permuted distribution (p=0.1%), supporting a strong association between the *CMT2stop* variant and increased TE expression (**Fig. 4e**).

### *FBX5* downregulates mCG in long TEs

While mean CHH methylation at long TEs was lower than for most other populations, mCG and mCHG levels were higher in CVI (**Fig. 3a-c**). GWAS on mCG levels revealed one Bonferroni significant peak at the end of chromosome 2 that includes a nonsense variant that truncates the F-box Armadillo (Arm)-repeat gene *F-BOX PROTEIN 5* (*FBX5*) (also known as *ARABIDILLO-1*) in the 4^th^ exon (Chr2:18,513,626, A>T, p.K555*) (**Fig. 5a, c**). *FBX5* is part of a E3 ubiquitin ligase complex (SCF) that was shown to regulate lateral root development and abscisic acid-mediated inhibition of seed germination in Arabidopsis^57–60^. *FBX5* has not previously been implicated in DNA methylation.

**Fig. 5.**
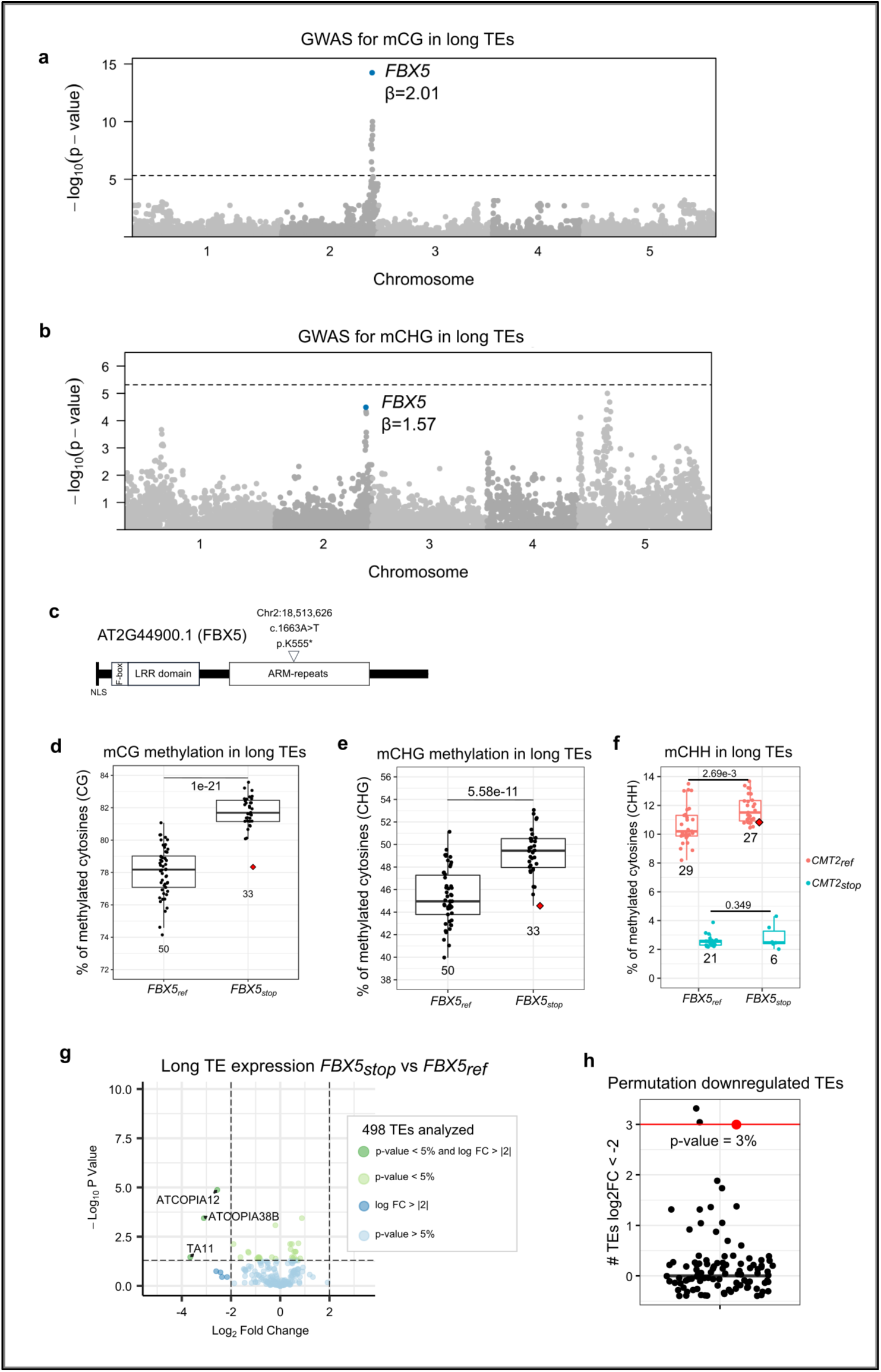
*FBX5* premature stop codon is associated with DNA methylation variation in CVI. Manhattan plots of the GWAS for mCG (**a**) and mCHG (**b**) in long TEs (>4kb). Nonsense SNP at *FBX5* is highlighted in blue. A dashed line indicates the Bonferroni threshold**. c,** Representation of the protein domains of FBX5*. W*hite triangle indicates the location of the nonsense mutation with HGVS nomenclature information. White boxes represent the protein domains (NLS: Nuclear localization signal, LRR: Leucine-rich repeat, ARM: ARMADILLO) as defined in previous work ^58^. Levels of mCG **(d)**, mCHG **(e)** and mCHH **(f)** at long TEs for accessions with the reference (*FBX5_ref_*) alternative (*FBX5_stop_*). For **f**, samples are separated based on their CMT2 allelic state. Center lines of boxplots show the medians, box limits show the 25th and 75th percentiles, whiskers extend 1.5 times the interquartile range from the 25th and 75th percentiles, and dots represent individual accessions. The number of individuals for each group is indicated below each box. Red diamonds represent the accession Cvi-0. Two-tailed Welch’s t-tests were performed to compare the means of the two groups in **d** and two-tailed Student’s t-tests were performed in **e** and **f**. P-values are indicated in the plots. **g**, Volcano plot of long TE expression in accessions with the *FBX5_stop_* vs *FBX5_ref_* alleles. The name of the TEs that have significant differential expression (< 5% adjusted p-value). **h**, Number of significantly downregulated TEs (log2FC < -2) in permutated alleles within populations for 100 permutations (black dots) and the observed value indicated in red (3 TEs). P-value is indicated on the plot and calculated by using the formula (B+1)/(M+1), where B is the number of permutations in which a value greater or equal than the observed value is obtained and M is the total number of permutations sampled.

There were no Bonferroni-significant peaks for mCHG in long TEs. However, a peak at *FBX5* that includes the p.K555* variant was the one of the strongest signals (**Fig. 5b**). The strongest peak for mCHG at long TEs was a complex peak on chromosome 5 (**Fig. 5b**) within 500 kb of three genes known to be involved in DNA methylation: *AGO9, DRM1*, and *SUVH4*. A peak around *MET1* was also visible at several Mb away at the distal end of chromosome 5. Of the four candidate genes we identified two contained coding variants with potential functions: a missense variant in *SUVH4*, and two missense variants in *MET1*.

Accessions carrying *FBX5_stop_* had 4.9% higher mCG (β = 2) and 6.3% higher mCHG levels (β = 1.57) (**Fig. 5d-e**). Although the *FBX5* region did not contain an obvious peak in the GWAS for mCHH (**Fig. 4a**), there was a moderate peak in the region. We examined the pattern within the CVI population and found evidence for a significant association of increased mCHH with *FBX5stop* within the CMT2ref background (**Fig. 5f**), suggesting that *CMT2* is epistatic to *FBX5* in the regulation of mCHH. Although the effect of FBX5 is clearest in the CG context, trends in our data and another recent publication (Zhang et al. co-submission), where peaks for mCHH and mCHG were identified in the same region in the Cvi-0 x Ler-0 RIL population, suggest that that FBX5 likely exerts a more general effect on DNA methylation.

*FBX5_stop_* is present in 13 populations with an overall frequency of 33% (62 out 189 accessions) in Santo Antão. We did not find other significant peaks when *FBX5_stop_* was used as a covariate. Surprisingly, although Cvi-0 is homozygous for the *FBX5_ref_* allele, it is at the extreme low end of the range of mCG and mCHG (**Fig. 5d-e**). This suggests that novel mutations may have arisen during the many successive propagations in artificial conditions since its initial collection 40 years ago in CVI and that some aspects of the Cvi-0 epigenome may not be representative of the natural population.

Converse to the pattern for CMT2*stop*, the *FBX5stop* allele was associated with increased DNA methylation over long TEs, in line with a de-repressive function of FBX5 reflected in the increased mCG and mCHG levels in the mutant (**Fig. 5g**). We found 30 significantly differentially expressed TEs between the two allelic states in the CVI population, including 16 downregulated and 14 upregulated TEs (**Fig. 5g**). The downregulated TEs belong to several superfamilies, including DNA/MuDR and LTR/Copia elements. AT2TE22635 (TA11 family, Line/L1 superfamily) showed the strongest downregulation in the FBX_stop_ background (**Fig. 5g**). Significantly upregulated TEs were all from the LTR/Gypsy superfamily, with 10 ATHILA and 4 ATLANTYS3 family TEs. We tested whether this pattern was likely to be found by chance using permutation analysis wherein we randomly attributed *FBX5* alleles within resampled individuals. We found that the number of downregulated TEs is greater than that ever found in 100 permutations (p=0.01), supporting that the *FBX5* variant is associated with significant downregulation of TE expression (**Fig. 5h**).

To confirm the role of *FBX5* in DNA methylation, we performed whole-genome bisulfite sequencing in two independent *fbx5* T-DNA insertion lines (SAIL_190_D02, SALK_082977), a double mutant of *fbx5* and its close homolog *arabidillo*-2 (SAIL_190_D02/SAIL_162_B11), and a transgenic *FBX5* overexpression line (FBX5-OE)^57^ (**Fig. 6a**). We found increased methylation levels for mCG (**Fig. 6b**) and mCHG in the T-DNA insertion lines. Consistent with this general pattern, mCG was lower, although not significantly reduced, in the *FBX5* overexpression line (**Fig. 6b**). The homologue *ARABIDILLO-2* also appears to impact mCG based on the higher mCG level in the double mutant compared to the single mutant (**Fig. 6b**). We also generated CRISPR *fbx5* knock-out lines in a CVI genetic background using the CVI accession S7-B5 that contained *CMT2_ref_* and *FBX5_ref_* (**Fig. 6a**). We produced two lines that were homozygous for the Cas9-induced mutations (*fbx5-1* and *fbx5-2*) and one heterozygous line (*fbx5-3*). Homozygous lines show an increase in long TE mCG, while the heterozygous line shows an intermediate phenotype, indicating haplo-insufficiency (**Fig. 6c**). No significant methylation changes were observed overall for CHG and CHH contexts, indicating that any impact in these contexts may change more subtly and stochastically over generations. These results confirm that *FBX5* is a negative regulator of DNA methylation, most strongly in the CG context.

**Fig. 6.**
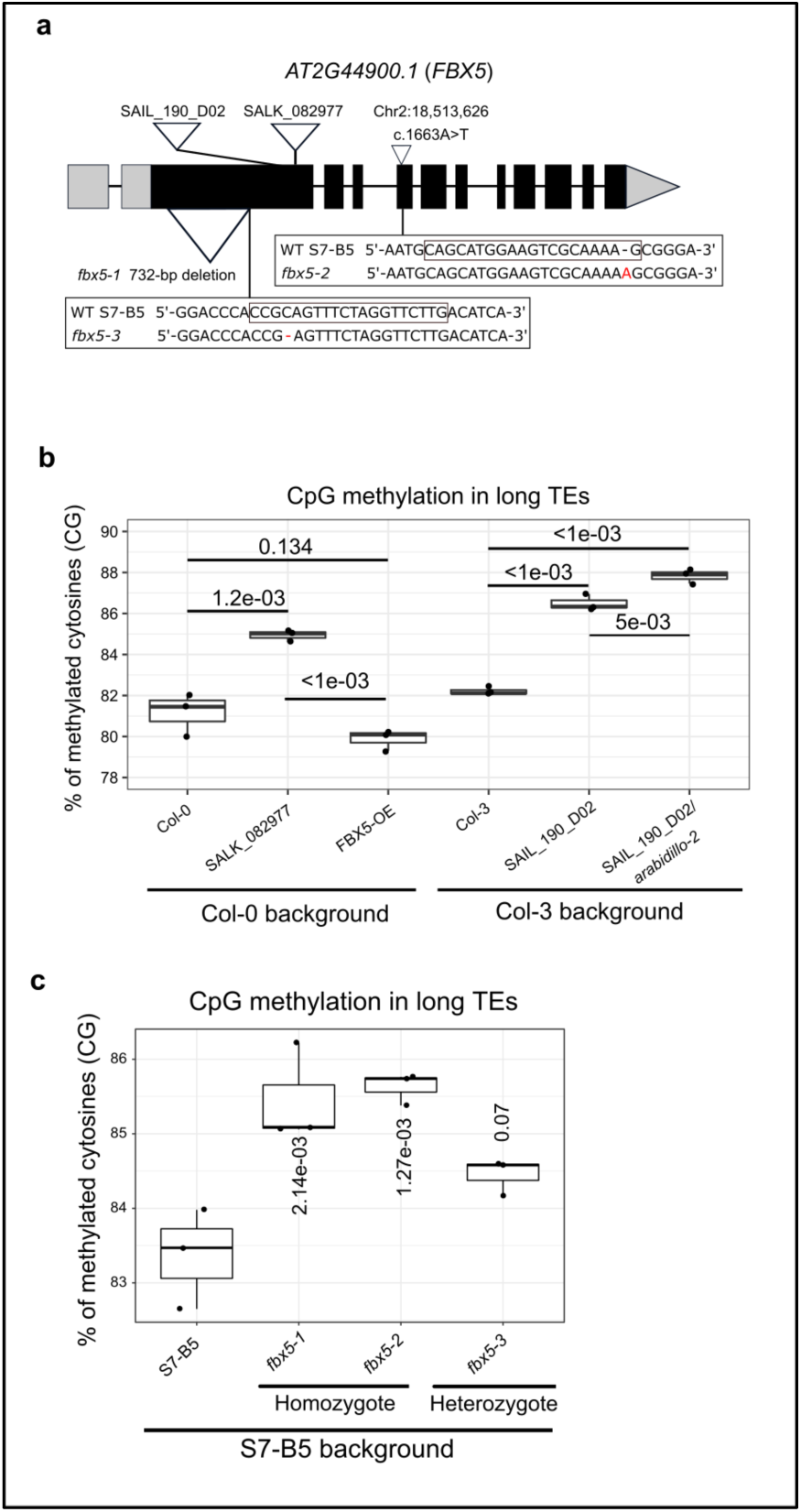
*FBX5* is a negative regulator of DNA methylation in long TEs. **a,** Representation of the transcript model *AT2G44900.1. W*hite triangle indicates the location of the nonsense mutation with HGVS nomenclature information. Grey boxes represent UTRs and black boxes the exons. T-DNA insertion lines are represented by white triangles with their stock names. The genotype of the three CRISPR/Cas-9 based frameshift mutants (*fbx5-1*, *fbx5-2*, and *fbx5-3*) are indicated in comparison to the background CPV accession used (S7-B5). The line *fbx5-1* shows a deletion of 720-bp (white triangle). The box in the wild-type S7-B5 sequence indicates the target site for CRISPR/Cas9, and red characters in mutants show the indel mutations. **b,** Level of mCG in long TEs for FBX5 T-DNA insertion lines SALK_082977, SAIL_190_D02, SAIL_190_D02/arabidillo-2 (SAIL_162_B11), overexpression line (FBX5-OE), and the background accessions used for transformation (Col-0 for SALK and Col-3 for SAIL). For each background, an ANOVA was performed followed by a Tukey’s test (two-tailed). P-values are indicated on the plot. **c,** Level of mCG in long TEs for S7-B5 (*FBX5_ref_*) and three independent *fbx5* CRISPR-Cas9 lines in the S7-B5 background (three biological replicates per line). The line *fbx5-3* is heterozygous for the mutation and shows an intermediate phenotype. An ANOVA was performed followed by a Dunnett’s test (two-tailed) with S7-B5 used as control. P-values are indicated in the plot. The center lines of boxplots show the medians, box limits show the 25th and 75th percentiles, whiskers extend 1.5 times the interquartile range from the 25th and 75th percentiles, and dots represent individual accessions.

## Discussion

Mutations that impact DNA methylation machinery have the potential to produce rapid genome-wide changes due to their widespread genomic impacts^61,62^. Our GWAS in CVI revealed high impact genetic polymorphisms for two known players of DNA methylation, *VIM2/4* and *CMT2*, and a novel player, *FBX5*.

A 2.7 kb deletion between the two *VARIANT IN METHYLATION* homologs *VIM2* and *VIM4* reduced gbM in CVI. In Arabidopsis, there are five *VIM* homologs (*VIM1–5*)^52,63^, which regulate DNA methylation by recruiting MET1 to hemi-methylated CG sites for maintenance methylation. This is accomplished with a PHD domain, which enables reading of core histones and an SRA domain, which facilitates binding to methylated DNA at histone H3 (reviewed in ^64^). The triple mutant *vim1/2/3* phenocopies the *met1* null allele mutant through complete loss of mCG^54,55^. Whereas a family of *VIMs* encode proteins required for methylation regulation in plants, in mammals a single orthologue, *UHRF1*, encodes a protein responsible for these functions^65,66^. Expansion of this gene family likely facilitated separation of functions and higher specificity. In the CVI population, we found differences in gene body methylation between*VIM2/4* alleles was strongly enriched for Cul4-RING E3 ubiquitin ligase complex, which include genes involved in environmental response traits, including light signaling, flowering, metabolism and DNA repair. The mammalian homolog of the VIMs, *UHRF1*, is involved in DNA repair^67^, suggesting that VIMs could also have such functions in plants. Overall, our results for the *VIM2/4* region point to additional hypotheses that could be tested related to the role of gene body methylation in stress response evolution.

We also identified genetic variants that impacted methylation levels over TEs. mCHH levels on TEs, especially long TEs, showed a bimodal distribution, which is due to an allele that disrupts *CMT2* that segregates in CVI. Other *CMT2* null alleles have been identified worldwide and their distribution is correlated with climate, suggesting that *CMT2* natural variation could be adaptive^44,68,69^. Further, two studies have shown that *cmt2* mutants are more resistant to heat stress and one study demonstrated a higher resistance to UV-B radiation^44,56^. However, disruption of CMT2 is also expected to reduce silencing of TEs, resulting in a potential increase in genome instability. A previous study found that COPIA transposon expression increased following heat stress in Col-0-*cmt2* and in the Kyoto accession, which carries a natural CMT2 mutation^70,71^. Since CMT2 is involved in maintenance methylation, its disruption likely does not immediately remove methylation marks; instead release of COPIA elements may be aided by heat exposure over generational time^72,73^. Consistent with this, we found that several COPIA elements were expressed in the *CMT2* null background in the CVI natural population, which is exposed to heat and drought stress conditions. The *CMT2stop* allele is segregating at intermediate frequencies across nearly all CVI sub-populations we sampled, suggesting they may be evolving under balancing selection where the allele provides a positive impact in resistance to heat and UV stress, but a negative impact in genome stability. An alternative possibility derives from the argument that the increased mutation rate due to TEs may be beneficial in a population adapting to a novel, harsh environment^72,74,75^. Under this logic, the increased mutation rate itself may have been to some degree beneficial in the variation-limited CVI population.

We discovered a novel player in DNA methylation, *FBX5* (*ARABIDILLO-1*), that impacts CG methylation over TEs. Similar to the *CMT2stop* variant in CVI, the *FBX5stop* allele is segregating in almost all subpopulations. The mild increase of mCG in accessions with the *FBX5* null allele was associated with the downregulation of several TEs, including COPIA elements, indicating that even a mild increase in DNA methylation can potentially lead to changes in TE repression. *FBX5* and its close homologue *ARABIDILLO-2* were previously shown to regulate lateral root development and abscisic acid-mediated inhibition of seed germination^57–60^. This indicates a pleiotropic effect of *FBX5* and *ARABIDILLO-2* on plant molecular and organismal phenotypes with potential adaptive consequences. Interestingly, the mammalian orthologue of FBX5, β-catenin, interacts physically with the mammalian orthologue of MET1, DNMT1, assuring mutual stabilization of the two proteins^76^. Considering that protein sequence similarity between FBX5 and β-catenin is high^77^, disruption of FBX5 as found in CVI could potentially affect MET1 activity. Further investigation is needed to determine how *FBX5* regulates DNA methylation.

While mutations that shape DNA methylation variation in CVI have major effects—disrupting DNA methylation regulators (*CMT2 K1166\**, *FBX5 K555\**) or removing a large shared regulatory region (*VIM2/4*)—the genes involved tend to impact DNA methylation in subtle ways. CMT2 functions in maintenance of DNA methylation so that its absence may act to quantitatively reduce rather than eliminate DNA methylation over generations. FBX5 likely interacts with MET1 based on studies of the human homologue, β-catenin^76^. However, we don’t observe DNA methylation phenotypes associated with major effect mutations in key DNA methylation genes like *DDM1* or *MET1*. This suggests natural variation may be more readily maintained in nature when it produces subtle effects on DNA methylation, allowing for tuning of DNA methylation rather than the large-scale loss that would result from major effect mutations in *MET1* or *DDM1*. Overall, our findings point to several potential areas for further investigation to dissect the roles of DNA methylation in environmental response.

## Material and methods

### Growth conditions

Plants were grown on soil (Brill Classic, Brill Substrate GmbH & Co. KG) and stratified one week at 4°C. Pots were transferred in a Bronson chamber (Bronson Climate BV) with 12h day/12h night photoperiod, light intensity 200 µmol/m^2^/s, temperature 21°C day/14°C night.

### Plant material

Seeds of accessions from Cape Verde and Morocco are available in our seed stock (Max Planck Institute for Plant Breeding Research, Cologne, Germany)^48,78^. Seeds of *fbx5* T-DNA mutants from Syngenta Arabidopsis Insertion Library (SAIL_190_D02) and SALK (SALK_082977), *arabidillo-2* (SAIL_162_B11), double mutant *fbx5/arabidillo-2* (SAIL_190_D02/SAIL_162_B11), and *FBX5* overexpressing lines were provided by Juliet Coates^57^. Seeds from the 1001GP accessions Dör-10 and UKID116, the *cmt2* T-DNA mutant (SAIL_906_G03), and Col-0 (CS76114) were ordered from the Nottingham Arabidopsis Stock Centre (NASC). T-DNA insertions were genotyped by PCR.

### DNA sequencing

Raw sequencing read data from whole-genome sequencing of the 190 Cape Verde (Santo Antão) accessions originate from published work^48^ and are available on European Nucleotide Archive (ENA) under accession code PRJEB39079.

### GWAS analysis

SNP calling was performed using MPI-SHORE pipeline^79^ with minor modifications. Detailed script is available at https://github.com/HancockLab/SNP_calling_Arabidopsis.

VCF processing and GWAS analysis was performed as detailed in repository https://github.com/johanzi/gwas_gemma. Briefly, non-biallelic, singletons, and indel variants were filtered out of the VCF file before analysis. Only SNP calls with a read coverage DP>=3 and genotype quality GQ>=25 were considered. To characterize the genetic architecture of the different traits, we used a Bayesian sparse linear mixed model (BSLMM) in GEMMA (version 0.94) (Zhou et al., 2013). We ran Markov chain Monte Carlo (MCMC) with 10,000,000 sampling steps and 2,500,000 burn-in iterations. We calculated the median and 95% confidence interval for the proportion of variance in phenotypes explained by available genotypes (PVE) across ten runs. A minor allele frequency (MAF) of 5% was applied so that the final set 10,321 analyzed SNPs were used for GWAS. A univariate LMM was used for association testing with the software GEMMA ^80^ according to the model:

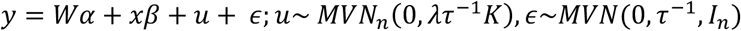

where **y** is an *n*-vector of quantitative traits (weighted DNA methylation level) for *n* individuals; **W**=(**w**_1_*; · · ·;* **w***_c_*) is an *n×c* matrix of covariates (fixed effects) including a column of 1s; ***α*** is a *c*-vector of the corresponding coefficients including the intercept; **x** is an *n*-vector of marker genotypes; *β* is the effect size of the marker; **u** is an *n*-vector of random effects; *ε* is an *n*-vector of errors; *τ_-_*_1_ is the variance of the residual errors; *λ* is the ratio between the two variance components; **K** is a known *n × n* relatedness matrix and **I***_n_* is an *n × n* identity matrix. MVN*_n_* denotes the *n*-dimensional multivariate normal distribution. We assessed SNP significance using the Bonferroni threshold (-log_10_(0.05/number of SNPs tested)). The standardized kinship matrix was used.

To integrate the deletion status at *VIM2/4* in the VCF file, which contains only SNPs, an artificial SNP coding for the absence/presence of the deletion was added manually at the approximate position of the deletion.

Beta (β) coefficients for the different SNPs of interest were obtained from the LMM of the GWAS and provided the effect size of each SNP on the phenotype. β coefficients are in the unit of the phenotype analyzed and should be interpreted as a slope coefficient where the value of the phenotype changes for each unit copy of the allele present, so the β value should be doubled when comparing diploid homozygous individuals. For instance, β=2 for SNP_x_ indicates an increase of four units of phenotype for the individual with the alternative allele of SNP_x_ in the homozygous state.

The *k*-mer GWAS was conducted as previously described^51^. The top 5% significant *k*-mers were mapped with bowtie 2 (version 2.4.4)^81^ to the four Santo Antão assembled genomes (see section below).

### Whole genome assemblies

Four accessions from Santo Antão (Cvi-0, S1-1, S5-10, and S15-3) were sequenced and assembled using long read sequencing by the by the Max Planck Genome Center in Cologne (Germany). Briefly, the genomic DNA was extracted from pooled leaf tissue from the same genotype. DNA fragments above 30 kb were selected with BluePippin (Sage Science). The libraries were prepared using Ligation Sequencing Kit 1D (Oxford Nanopore Technologies, catalog no. SQK-LSK109) and sequenced using the GridION X5 platform (Oxford Nanopore Technologies). De novo assemblies were generated with miniasm-0.3^82^ and minimap2-2.17^83^. The resulting draft assemblies were first polished twice with racon (version 1.4.10)^84^ using the raw Nanopore reads and then 10 more times with pilon (version 1.23)^85^ using the corresponding Illumina short reads. We oriented and scaffolded the contigs using the reference genome assembly (TAIR10). To prevent mapping of Illumina short reads to anchoring points of the contigs, we inserted 1000 N’s between the contigs. The pipeline can be found in https://github.com/HancockLab/Fogo-Edaphic. Nanopore reads mapping the *VIM2/4* region are present in NCBI under accession code PRJNA1112558.

### Whole genome bisulfite sequencing

Rosette leaves were harvested 20 to 25 days after germination (DAG) at stage 4-8 true leaves. Collect was performed between Zeitgeber (ZT) 3 and 6 for whole-genome bisulfite sequencing. Samples were directly flash-frozen in liquid nitrogen after collection. Whole-genome bisulfite sequencing libraries were prepared as described in^86^. Libraries were sequenced on HiSeq3000 in 150 bp single-end mode to generate about 7 million reads of data per library. The adaptors were trimmed using Cutadapt (v2.9) ^87^ and mapped on TAIR10 reference genome with Bismark (v0.19.0) (parameters –N 1)^88^. The conversion efficiency was estimated using the levels of unconverted cytosines found in the chloroplast unmethylated genome. Levels of converted cytosines were higher than 99% in all cases. Cytosine reports from Bismark were imported in R and subsequent methylation analyses were performed with the methylKit package (v1.14.2)^89^. Differentially methylated regions (DMRs) were defined with window size of 300 bp and an overlapping of window of 100 bp. Overlapping DMRs with a methylation difference > 25% and a q-value < 0.01 were merged and their differential methylation levels were averaged.

To compared our data to the 1001GP analysis, we bisulfite-sequenced the two outlier accessions Dör-10 (seqID 5856, Sweden) and UKID116 (seqID 5822, the United Kingdom), which showed high and low gbM, respectively^45^ and Col-0, which exhibits an intermediate value. Levels of gbM for UKID116 and Dör-10 in our data diverge from only 1-2% of the values from the 1001GP data using our analysis pipeline. The gbM levels of these two accessions encompasses the values of gbM found of the Moroccan accessions, with Cvi-0 being close to UKID116 and Col-0 showing intermediate values, as previously reported.

### Whole genome transcriptome sequencing

We generated leaf transcriptomes for 97 Santo Antão accessions, including 11 accessions with three biological replicates. Rosette leaves from 4-true leaves (20 DAG) were collected between ZT3 and ZT6 and flash-frozen into liquid nitrogen. Samples were ground in 2 ml Eppendorf tubes containing one tungsten carbide ball in a TissueLyser II (Qiagen). An aliquot of about 20 mg of powder was transferred into a 96 well plate, and total RNA was prepared with the NucleoMag® 96 RNA kit (Macherey Nagel). Libraries were prepared with the NEBNext Ultra™ Directional RNA Library Prep Kit for Illumina sequencing (New England Biolabs). Approximately about 7 million reads of 150 bp single-end reads were generated on the Illumina sequencer HiSeq3000. The adaptors were trimmed using Cutadapt (v3.5) (parameters -m 20 -q 35)^87^. Reads were mapped on TAIR10 reference genome and Araport11 gene annotation using HISAT2 (v2.2.0)^90^. The read count was performed with HTSeq (v0.12.4)^91^. Differential expression analysis was performed in R using the DESeq2 package (v1.28.1)^92^. Biological replicates of the eleven accessions cluster tightly on the principal component analysis (PCA) for gene expression, indicating that environmental effects on gbM within the growth chamber were minimum.

### VIM2/4 deletion analysis

To detect the *VIM2/VIM4* deletion in the Cape Verde accessions sequenced, we could not rely on the broken paired-end reads due to the high homology between *VIM2* and *VIM4*. Instead, we defined a minimum sequencing coverage at the defined location of the deletion below which we consider an accession as having the *VIM2/VIM4* deletion. Based on the read distribution at the deletion site (chr1:24586731-24589471) across the sequenced accessions, we defined accessions as having the *VIM2/4* deletion when the number of reads mapping at the deletion was below 50. The read count was performed with Samtools (v1.9).

### Production and analysis of CRISPR lines

CRISPR-Cas9 transgenic lines for *FBX5* were generated as previously described ^93^ with details available in https://github.com/johanzi/crispr_cas9_arabidopsis. Transformed T1 plants in the S7-B5 background (CVI accession with *FBX5_ref_* allele) were recovered from hygromycin plates and transplanted in soil. Mutations at the target sites were verified using Sanger sequencing. We also generated high-coverage (ranging from 59.7 to 81.1x) short-read genomic data for the CRISPR lines to investigate the potential for off-target mutations. We extracted DNA with the DNeasy Plant Mini Kit (Qiagen, catalog no. 69106) and prepared the libraries using NEBNext Ultra II FS DNA Library Prep [Illumina, New England Biolabs (NEB)]. Sequencing was conducted on an Illumina HiSeq3000 platform (Illumina, NEB) with 68-82 million reads (2x150 bp) (Novogene). We examined potential off-target mutations as described previously^94^. Briefly, the resulting reads were mapped to TAIR10 with bwa (version 0.7.15) using the mem algorithm. We used GATK to call SNPs and short indels. We predicted potential off-target sites with Cas-OFFinder^95^. We used a maximum of five mismatches, SpCas9 from Streptococcus pyogenes (5′-NGG-3′) as CRISPR-Cas-derived RNA-guided endonuclease and TAIR10 as a target reference genome. Finally, we examined evidence for overlap between predicted off-target sites and SNPs and indels with a custom python script (https://github.com/HancockLab/Fogo-Edaphic). We found no evidence of off-target mutations in the CRISPR lines. T3 seeds of three independent *FBX5* CRISPR-Cas9 transgenic lines were used for WGBS.

## Data availability

Raw sequencing read data for whole-genome bisulfite sequencing and RNA-seq were deposited in NCBI under the accession code PRJNA612437. The Nanopore whole-genome assemblies of the SA accessions S1-1, S15-3, S5-10 and Cvi-0 (region Chr1-24,400,000-24,800,000) are available in PRJNA1112558. Whole-genome sequencing of CRISPR lines for off-target analysis are available in PRJNA1112441.

## Code availability

Scripts of the bioinformatics analyses are available on GitHub (https://github.com/johanzi/scripts_cvi_methylation_variants).

## Acknowledgments

This work was supported by the ERC starting grant CVI_ADAPT 638810 and core funding from the Max Planck Society to A.M.H. and by the National Science Foundation (MCB-2242696) to R.J.S. Paula Unger and Nina Doering helped with the experiments. We thank Damon Lisch for helpful comments on the manuscript and Eric Richards for discussion regarding analysis of the VIM regions.

## Author contributions

Conceptualization, A.M.H. and J.Z.; Methodology, A.M.H., R.J.S and J.Z.; Investigation, A.M.H., J.Z., E.T., A.F.; Writing –Original Draft, A.M.H. and J.Z.; Writing –Review & Editing, A.M.H., J.Z., and R.J.S.; Funding Acquisition, A.M.H.; Resources, E.M., C.N., A.E.; Supervision, A.M.H. and J.Z.; M.G.: SNP GATK pipeline, long read assembly; C.N.: Material collection and propagation. M.G.: Long-read assemblies of the four CVI accessions used for *VIM2/4* deletion characterization.

## References

1. Law, J. A. & Jacobsen, S. E. Establishing, maintaining and modifying DNA methylation patterns in plants and animals. Nat. Rev. Genet. 11, 204–220 (2010).

2. Jullien, P. E., Susaki, D., Yelagandula, R., Higashiyama, T. & Berger, F. DNA Methylation Dynamics during Sexual Reproduction in *Arabidopsis thaliana*. Curr. Biol. 22, 1825–1830 (2012).

3. Pikaard, C. S. & Scheid, O. M. Epigenetic Regulation in Plants. Cold Spring Harb. Perspect. Biol. 6, a019315 (2014).

4. Zhang, H., Lang, Z. & Zhu, J.-K. Dynamics and function of DNA methylation in plants. Nat. Rev. Mol. Cell Biol. 19, 489–506 (2018).

5. Bond, D. M. & Baulcombe, D. C. Small RNAs and heritable epigenetic variation in plants. Trends Cell Biol. 24, 100–107 (2014).

6. Jean Finnegan, E. & Dennis, E. S. Isolation and identification by sequence homology of a putative cytosine methyltransferase from Arabidopsis thaliana. Nucleic Acids Res. 21, 2383– 2388 (1993).

7. Finnegan, E. J., Peacock, W. J. & Dennis, E. S. Reduced DNA methylation in Arabidopsis thaliana results in abnormal plant development. Proc. Natl. Acad. Sci. 93, 8449–8454 (1996).

8. Bartee, L., Malagnac, F. & Bender, J. Arabidopsis cmt3 chromomethylase mutations block non-CG methylation and silencing of an endogenous gene. Genes Dev. 15, 1753–1758 (2001).

9. Lindroth, A. M. et al. Requirement of CHROMOMETHYLASE3 for Maintenance of CpXpG Methylation. Science 292, 2077–2080 (2001).

10. Kankel, M. W. et al. Arabidopsis MET1 cytosine methyltransferase mutants. Genetics 163, 1109–1122 (2003).

11. Du, J. et al. Dual Binding of Chromomethylase Domains to H3K9me2-Containing Nucleosomes Directs DNA Methylation in Plants. Cell 151, 167–180 (2012).

12. Stroud, H. et al. Non-CG methylation patterns shape the epigenetic landscape in Arabidopsis. Nat. Struct. Mol. Biol. 21, 64–72 (2014).

13. Cao, X. et al. Conserved plant genes with similarity to mammalian de novo DNA methyltransferases. Proc. Natl. Acad. Sci. 97, 4979–4984 (2000).

14. Matzke, M. A. & Mosher, R. A. RNA-directed DNA methylation: an epigenetic pathway of increasing complexity. Nat. Rev. Genet. 15, 394 (2014).

15. Zemach, A. et al. The Arabidopsis Nucleosome Remodeler DDM1 Allows DNA Methyltransferases to Access H1-Containing Heterochromatin. Cell 153, 193–205 (2013).

16. Jackson, J. P., Lindroth, A. M., Cao, X. & Jacobsen, S. E. Control of CpNpG DNA methylation by the KRYPTONITE histone H3 methyltransferase. Nature 416, 556–560 (2002).

17. Malagnac, F., Bartee, L. & Bender, J. An Arabidopsis SET domain protein required for maintenance but not establishment of DNA methylation. EMBO J. 21, 6842–6852 (2002).

18. Tran, R. K. et al. Chromatin and siRNA pathways cooperate to maintain DNA methylation of small transposable elements in Arabidopsis. Genome Biol. 6, R90 (2005).

19. Du, J. et al. Mechanism of DNA Methylation-Directed Histone Methylation by KRYPTONITE. Mol. Cell 55, 495–504 (2014).

20. Johannes, F. et al. Assessing the Impact of Transgenerational Epigenetic Variation on Complex Traits. PLOS Genet. 5, e1000530 (2009).

21. He, L. et al. DNA methylation-free Arabidopsis reveals crucial roles of DNA methylation in regulating gene expression and development. Nat. Commun. 13, 1335 (2022).

22. Deaton, A. M. & Bird, A. CpG islands and the regulation of transcription. Genes Dev. 25, 1010–1022 (2011).

23. Schmitz, R. J., Lewis, Z. A. & Goll, M. G. DNA Methylation: Shared and Divergent Features across Eukaryotes. Trends Genet. 35, 818–827 (2019).

24. Tran, R. K. et al. DNA Methylation Profiling Identifies CG Methylation Clusters in Arabidopsis Genes. Curr. Biol. 15, 154–159 (2005).

25. Zhang, X. et al. Genome-wide High-Resolution Mapping and Functional Analysis of DNA Methylation in Arabidopsis. Cell 126, 1189–1201 (2006).

26. Cokus, S. J. et al. Shotgun bisulphite sequencing of the Arabidopsis genome reveals DNA methylation patterning. Nature 452, 215–219 (2008).

27. Lister, R. et al. Highly Integrated Single-Base Resolution Maps of the Epigenome in Arabidopsis. Cell 133, 523–536 (2008).

28. Takuno, S. & Gaut, B. S. Body-Methylated Genes in Arabidopsis thaliana Are Functionally Important and Evolve Slowly. Mol. Biol. Evol. 29, 219–227 (2012).

29. Takuno, S. & Gaut, B. S. Gene body methylation is conserved between plant orthologs and is of evolutionary consequence. Proc. Natl. Acad. Sci. 110, 1797–1802 (2013).

30. Niederhuth, C. E. et al. Widespread natural variation of DNA methylation within angiosperms. Genome Biol. 17, 194 (2016).

31. Vaughn, M. W. et al. Epigenetic Natural Variation in Arabidopsis thaliana. PLOS Biol 5, e174 (2007).

32. Zilberman, D., Gehring, M., Tran, R. K., Ballinger, T. & Henikoff, S. Genome-wide analysis of *Arabidopsis thaliana* DNA methylation uncovers an interdependence between methylation and transcription. Nat. Genet. 39, 61–69 (2007).

33. Saze, H. & Kakutani, T. Differentiation of epigenetic modifications between transposons and genes. Curr. Opin. Plant Biol. 14, 81–87 (2011).

34. Horvath, R., Laenen, B., Takuno, S. & Slotte, T. Single-cell expression noise and gene-body methylation in Arabidopsis thaliana. Heredity 123, 81–91 (2019).

35. Bewick, A. J. et al. On the origin and evolutionary consequences of gene body DNA methylation. Proc. Natl. Acad. Sci. 113, 9111–9116 (2016).

36. Kiefer, C. et al. Interspecies association mapping links reduced CG to TG substitution rates to the loss of gene-body methylation. Nat. Plants 5, 846–855 (2019).

37. Bewick, A. J. & Schmitz, R. J. Gene body DNA methylation in plants. Curr. Opin. Plant Biol. 36, 103–110 (2017).

38. Zilberman, D. An evolutionary case for functional gene body methylation in plants and animals. Genome Biol. 18, 87 (2017).

39. Muyle, A. M., Seymour, D. K., Lv, Y., Huettel, B. & Gaut, B. S. Gene Body Methylation in Plants: Mechanisms, Functions, and Important Implications for Understanding Evolutionary Processes. Genome Biol. Evol. 14, evac038 (2022).

40. Williams, C. J., Dai, D., Tran, K. A., Monroe, J. G. & Williams, B. P. Dynamic DNA methylation turnover in gene bodies is associated with enhanced gene expression plasticity in plants. Genome Biol. 24, 227 (2023).

41. Coleman-Derr, D. & Zilberman, D. Deposition of Histone Variant H2A.Z within Gene Bodies Regulates Responsive Genes. PLOS Genet. 8, e1002988 (2012).

42. Muyle, A., Ross-Ibarra, J., Seymour, D. K. & Gaut, B. S. Gene body methylation is under selection in Arabidopsis thaliana. Genetics 218, iyab061 (2021).

43. Schmitz, R. J. et al. Patterns of population epigenomic diversity. Nature 495, 193–198 (2013).

44. Shen, X. et al. Natural CMT2 Variation Is Associated With Genome-Wide Methylation Changes and Temperature Seasonality. PLOS Genet. 10, e1004842 (2014).

45. Kawakatsu, T. et al. Epigenomic Diversity in a Global Collection of Arabidopsis thaliana Accessions. Cell 166, 492–505 (2016).

46. Pignatta, D. et al. Natural epigenetic polymorphisms lead to intraspecific variation in Arabidopsis gene imprinting. eLife 3, e03198 (2014).

47. Picard, C. L. & Gehring, M. Proximal methylation features associated with nonrandom changes in gene body methylation. Genome Biol. 18, 73 (2017).

48. Fulgione, A. et al. Parallel reduction in flowering time from de novo mutations enable evolutionary rescue in colonizing lineages. Nat. Commun. 13, 1461 (2022).

49. Yang, J., Zeng, J., Goddard, M. E., Wray, N. R. & Visscher, P. M. Concepts, estimation and interpretation of SNP-based heritability. Nat. Genet. 49, 1304–1310 (2017).

50. Zhou, X., Carbonetto, P. & Stephens, M. Polygenic Modeling with Bayesian Sparse Linear Mixed Models. PLOS Genet. 9, e1003264 (2013).

51. Voichek, Y. & Weigel, D. Identifying genetic variants underlying phenotypic variation in plants without complete genomes. Nat. Genet. 52, 534–540 (2020).

52. Woo, H. R., Pontes, O., Pikaard, C. S. & Richards, E. J. VIM1, a methylcytosine-binding protein required for centromeric heterochromatinization. Genes Dev. 21, 267–277 (2007).

53. Woo, H. R., Dittmer, T. A. & Richards, E. J. Three SRA-Domain Methylcytosine-Binding Proteins Cooperate to Maintain Global CpG Methylation and Epigenetic Silencing in Arabidopsis. PLOS Genet. 4, e1000156 (2008).

54. Feng, S. et al. Conservation and divergence of methylation patterning in plants and animals. Proc. Natl. Acad. Sci. 107, 8689–8694 (2010).

55. Stroud, H., Greenberg, M. V. C., Feng, S., Bernatavichute, Y. V. & Jacobsen, S. E. Comprehensive Analysis of Silencing Mutants Reveals Complex Regulation of the Arabidopsis Methylome. Cell 152, 352–364 (2013).

56. Jiang, J. et al. Substrate specificity and protein stability drive the divergence of plant-specific DNA methyltransferases. Sci. Adv. 10, eadr2222 (2024).

57. Coates, J. C., Laplaze, L. & Haseloff, J. Armadillo-related proteins promote lateral root development in Arabidopsis. Proc. Natl. Acad. Sci. 103, 1621–1626 (2006).

58. Nibau, C. et al. ARABIDILLO proteins have a novel and conserved domain structure important for the regulation of their stability. Plant Mol. Biol. 75, 77–92 (2011).

59. Gibbs, D. J. et al. AtMYB93 is a novel negative regulator of lateral root development in Arabidopsis. New Phytol. 203, 1194 (2014).

60. Moody, L. A. et al. An ancient and conserved function for Armadillo-related proteins in the control of spore and seed germination by abscisic acid. New Phytol. 211, 940–951 (2016).

61. Roux, F. et al. Genome-Wide Epigenetic Perturbation Jump-Starts Patterns of Heritable Variation Found in Nature. Genetics 188, 1015–1017 (2011).

62. Kooke, R. et al. Epigenetic Basis of Morphological Variation and Phenotypic Plasticity in Arabidopsis thaliana. Plant Cell 27, 337–348 (2015).

63. Johnson, L. M. et al. The SRA Methyl-Cytosine-Binding Domain Links DNA and Histone Methylation. Curr. Biol. 17, 379–384 (2007).

64. Grimanelli, D. & Ingouff, M. DNA Methylation Readers in Plants. J. Mol. Biol. (2020) doi:10.1016/j.jmb.2019.12.043.

65. Bostick, M. et al. UHRF1 Plays a Role in Maintaining DNA Methylation in Mammalian Cells. Science 317, 1760–1764 (2007).

66. Sharif, J. et al. The SRA protein Np95 mediates epigenetic inheritance by recruiting Dnmt1 to methylated DNA. Nature 450, 908–912 (2007).

67. Jin, B. & Robertson, K. D. DNA Methyltransferases (DNMTs), DNA Damage Repair, and Cancer. Adv. Exp. Med. Biol. 754, 3–29 (2013).

68. Dubin, M. J. et al. DNA methylation in Arabidopsis has a genetic basis and shows evidence of local adaptation. eLife 4, e05255 (2015).

69. Sasaki, E., Kawakatsu, T., Ecker, J. R. & Nordborg, M. Common alleles of CMT2 and NRPE1 are major determinants of CHH methylation variation in Arabidopsis thaliana. PLOS Genet. 15, e1008492 (2019).

70. Liu, S. et al. Role of H1 and DNA methylation in selective regulation of transposable elements during heat stress. New Phytol. 229, 2238–2250 (2021).

71. Nozawa, K. et al. Epigenetic regulation of ecotype-specific expression of the heat-activated transposon ONSEN. Front. Plant Sci. 13, 899105 (2022).

72. Pecinka, A. et al. Epigenetic Regulation of Repetitive Elements Is Attenuated by Prolonged Heat Stress in Arabidopsis. Plant Cell Online 22, 3118–3129 (2010).

73. Cavrak, V. V. et al. How a Retrotransposon Exploits the Plant’s Heat Stress Response for Its Activation. PLOS Genet 10, e1004115 (2014).

74. Dubin, M. J., Mittelsten Scheid, O. & Becker, C. Transposons: a blessing curse. Curr. Opin. Plant Biol. 42, 23–29 (2018).

75. Baduel, P. et al. Genetic and environmental modulation of transposition shapes the evolutionary potential of Arabidopsis thaliana. Genome Biol. 22, 138 (2021).

76. Song, J. et al. A Protein Interaction between β-Catenin and Dnmt1 Regulates Wnt Signaling and DNA Methylation in Colorectal Cancer Cells. Mol. Cancer Res. 13, 969–981 (2015).

77. Coates, J. C. Armadillo repeat proteins: beyond the animal kingdom. Trends Cell Biol. 13, 463–471 (2003).

78. Brennan, A. C. et al. The genetic structure of Arabidopsis thaliana in the south-western Mediterranean range reveals a shared history between North Africa and southern Europe. BMC Plant Biol. 14, 17 (2014).

79. Ossowski, S. et al. Sequencing of natural strains of Arabidopsis thaliana with short reads. Genome Res. 18, 2024–2033 (2008).

80. Zhou, X. & Stephens, M. Genome-wide efficient mixed-model analysis for association studies. Nat. Genet. 44, 821–824 (2012).

81. Langmead, B. & Salzberg, S. L. Fast gapped-read alignment with Bowtie 2. Nat. Methods 9, 357–359 (2012).

82. Li, H. Minimap and miniasm: fast mapping and de novo assembly for noisy long sequences. Bioinformatics 32, 2103–2110 (2016).

83. Li, H. Minimap2: pairwise alignment for nucleotide sequences. Bioinformatics 34, 3094–3100 (2018).

84. Vaser, R., Sović, I., Nagarajan, N. & Šikić, M. Fast and accurate de novo genome assembly from long uncorrected reads. Genome Res. 27, 737–746 (2017).

85. Walker, B. J. et al. Pilon: An Integrated Tool for Comprehensive Microbial Variant Detection and Genome Assembly Improvement. PLOS ONE 9, e112963 (2014).

86. Urich, M. A., Nery, J. R., Lister, R., Schmitz, R. J. & Ecker, J. R. MethylC-seq library preparation for base-resolution whole-genome bisulfite sequencing. Nat. Protoc. 10, 475–483 (2015).

87. Martin, M. Cutadapt removes adapter sequences from high-throughput sequencing reads. EMBnet.journal 17, 10–12 (2011).

88. Krueger, F. & Andrews, S. R. Bismark: a flexible aligner and methylation caller for Bisulfite-Seq applications. Bioinformatics 27, 1571–1572 (2011).

89. Akalin, A. et al. methylKit: a comprehensive R package for the analysis of genome-wide DNA methylation profiles. Genome Biol. 13, R87 (2012).

90. Kim, D., Paggi, J. M., Park, C., Bennett, C. & Salzberg, S. L. Graph-based genome alignment and genotyping with HISAT2 and HISAT-genotype. Nat. Biotechnol. 37, 907–915 (2019).

91. Anders, S., Pyl, P. T. & Huber, W. HTSeq—a Python framework to work with high-throughput sequencing data. Bioinformatics 31, 166–169 (2015).

92. Love, M. I., Huber, W. & Anders, S. Moderated estimation of fold change and dispersion for RNA-seq data with DESeq2. Genome Biol. 15, 550 (2014).

93. Wang, Z.-P. et al. Egg cell-specific promoter-controlled CRISPR/Cas9 efficiently generates homozygous mutants for multiple target genes in Arabidopsis in a single generation. Genome Biol. 16, 144 (2015).

94. Tergemina, E. et al. A two-step adaptive walk rewires nutrient transport in a challenging edaphic environment. Sci. Adv. 8, eabm9385 (2022).

95. Bae, S., Park, J. & Kim, J.-S. Cas-OFFinder: a fast and versatile algorithm that searches for potential off-target sites of Cas9 RNA-guided endonucleases. Bioinformatics 30, 1473–1475 (2014).

